# Movement ecology of captive-bred axolotls in restored and artificial wetlands: conservation insights for amphibian reintroductions and translocations

**DOI:** 10.1101/2024.11.09.622816

**Authors:** Alejandra G. Ramos, Horacio Mena, David Schneider, Luis Zambrano

## Abstract

Amphibians are among the most endangered vertebrates globally due to habitat loss, environmental degradation, and urban expansion. The Mexican axolotl (*Ambystoma mexicanum*), a critically endangered aquatic species endemic to Xochimilco, exemplifies these challenges. This study evaluates the viability of restored and artificial wetlands for axolotl conservation by comparing movement patterns, home range sizes, and habitat use. Using VHF telemetry, we tracked captive-bred axolotls released into both environments. Axolotls survived and foraged successfully in both sites, with those in La Cantera Oriente (LCO) exhibiting larger home ranges and greater daily distances traveled than those in Xochimilco. A quadratic relationship between water temperature and movement indicated a narrow thermal preference. Additionally, age and sex influenced movement patterns, with younger axolotls and females traveling greater distances. Recaptured individuals gained weight, suggesting successful adaptation, although two axolotls were lost to avian predation in Xochimilco after the study concluded. These findings highlight the potential of artificial wetlands like LCO for axolotl conservation by providing stable conditions that may mitigate habitat degradation and climate change impacts. The study recommends integrating native and artificial habitats into conservation strategies, incorporating predator awareness training before release, and ongoing habitat monitoring to enhance survival outcomes for this iconic species.

## Introduction

Amphibians are the most threatened group of vertebrates worldwide, with 41% of species currently listed as at risk of extinction on the International Union for Conservation of Nature (IUCN) Red List [1,2]. Additionally, approximately 25% of species are classified as Data Deficient, with estimates suggesting that half of these species are also likely threatened [3]. Amphibians play a crucial role in sustaining ecosystem dynamics as both predators and prey in wetlands and serve as essential indicators of biodiversity and environmental health [4–6]. Habitat loss, fragmentation, and degradation are among the primary drivers of global amphibian extinctions and population declines [7–10]. Urban expansion severely transforms freshwater ecosystems [5,11], disproportionately threatening amphibian species that are entirely dependent on aquatic habitats more than those that transition to terrestrial habitats as adults [12].

One such species is the critically endangered Mexican axolotl (*Ambystoma mexicanum*) [2], a paedomorphic salamander that retains its juvenile features and aquatic lifestyle into adulthood [13]. Endemic to Xochimilco, the last remnant of an extensive wetland system that once spanned the Central Valley of Mexico [14], the axolotl’s natural habitat has been significantly reduced and altered. Xochimilco has been managed for more than 1,500 years. Historically, during the Aztec Empire, the wetland was partially transformed into a series of canals surrounding rectangular islands called chinampas, which supported intensive food production and promoted local biodiversity [15]. However, urban expansion, environmental degradation, the introduction of invasive species, and a dramatic decrease in water quality due to surrounding urbanization have resulted in a marked decline in axolotl populations [16,17]. Restoration efforts, such as the ‘chinampa-refuge’ project initiated in 2004, aim to couple traditional food production with aquatic conservation [15]. Nevertheless, much of Xochimilco continues to face significant pressures from urban expansion and low water quality, limiting the effectiveness of these conservation efforts to the restored chinampas.

Given the intensifying amphibian extinction crisis, continuous and proactive human interventions are now considered essential to preserve vulnerable species and prevent extinctions [18–21]. Translocation—the movement of living organisms from one area to another—is an important conservation tool often combined with captive breeding to restore dwindling or extirpated populations and to establish new populations in suitable habitats [22,23]. Although most translocations have involved birds and mammals [19,24], amphibian translocations are increasing in number and success [22,25]. However, programs for threatened salamanders remain scarce, primarily because few species are easily kept and bred in captivity [26]. Concerns about the fitness of captive-reared animals and their ability to adapt to wild conditions persist, with predation-induced mortality being a main cause of translocation failure [27]. Additionally, not addressing the initial causes of population decline before releasing organisms can hinder the success of these efforts [28].

The creation and restoration of wetlands have proven effective for the conservation of numerous amphibian species, including salamanders [29]. For instance, the establishment of over 200 ponds led to increases in both amphibian diversity and populations of the threatened crested newt (*Triturus cristatus*) [30]. Understanding the specific habitat preferences, movement behaviors, and survival rates of threatened species is essential for designing effective conservation programs because distinct microhabitats within an ecosystem can differentially affect species fitness [31]. Previous studies on amphibians like hellbenders and Chinese giant salamanders have shown that factors such as age, sex, and habitat characteristics play critical roles in determining movement patterns [32,33]. Nevertheless, relatively few programs have examined post-release survival and habitat use in detail [26]. Biotelemetry methods, such as very high frequency (VHF) radio telemetry, offer valuable tools for monitoring individuals, allowing researchers to assess movement ecology, habitat use, and factors influencing post-release success [34].

While conserving threatened species within their natural habitats remains the ideal strategy [35], the ongoing challenges in Xochimilco encourage the exploration of alternative habitats for axolotl conservation. Furthermore, as climate unpredictability is expected to worsen, it becomes even more crucial to identify and protect refuges that are anticipated to remain suitable over time [6]. La Cantera Oriente (LCO) is an artificial wetland with four ponds and a stream, created around 30 years ago when basalt mining activities caused underground springs to surface. Despite its recent formation, LCO has developed into a biologically diverse habitat with a variety of aquatic vegetation [36]. Situated within the protected area of the Reserva Ecológica del Pedregal de San Ángel (REPSA), it is accessible primarily to students and researchers from the Universidad Nacional Autónoma de México (UNAM). The wetland supports native fauna, including potential prey and predators relevant to axolotls, and recent eradication efforts have reduced invasive species. Given these characteristics, LCO could function as a potential habitat for axolotls, offering an alternative site for conservation efforts outside their native environment.

In this study, we employed very high frequency (VHF) telemetry to evaluate the movement behaviors, home range sizes, and habitat use of captive-bred axolotls released into both an artificial wetland (LCO) and a restored chinampa within their native habitat of Xochimilco. By monitoring these individuals and analyzing factors influencing their movement and survival—such as environmental conditions and individual traits—we aim to assess the feasibility of both sites as viable locations for axolotl conservation. Our study offers practical implications for habitat restoration, reintroduction, and translocation programs, highlighting how both restored native habitats and artificial environments can play crucial roles in sustaining endangered species. Ultimately, this research aims to inform future conservation strategies by exploring how axolotls respond to different environmental conditions and emphasizing the importance of integrating both native and alternative habitats in their preservation.

## Materials and methods

### Study areas

This study was conducted in two distinct aquatic wetlands located in southern Mexico City: Xochimilco and La Cantera Oriente (LCO) (Fig 1). In Xochimilco, we used a 500 m² chinampa within the chinampa-refuge project, where canals are maintained without agrochemicals and equipped with filters to improve water quality and exclude exotic fish [37]. These refuges have proven beneficial for the survival and reproduction of axolotls and other native species [38]. They aim to recreate conditions akin to those in historical Xochimilco, where axolotl populations were once healthy.

**Figure 1.**
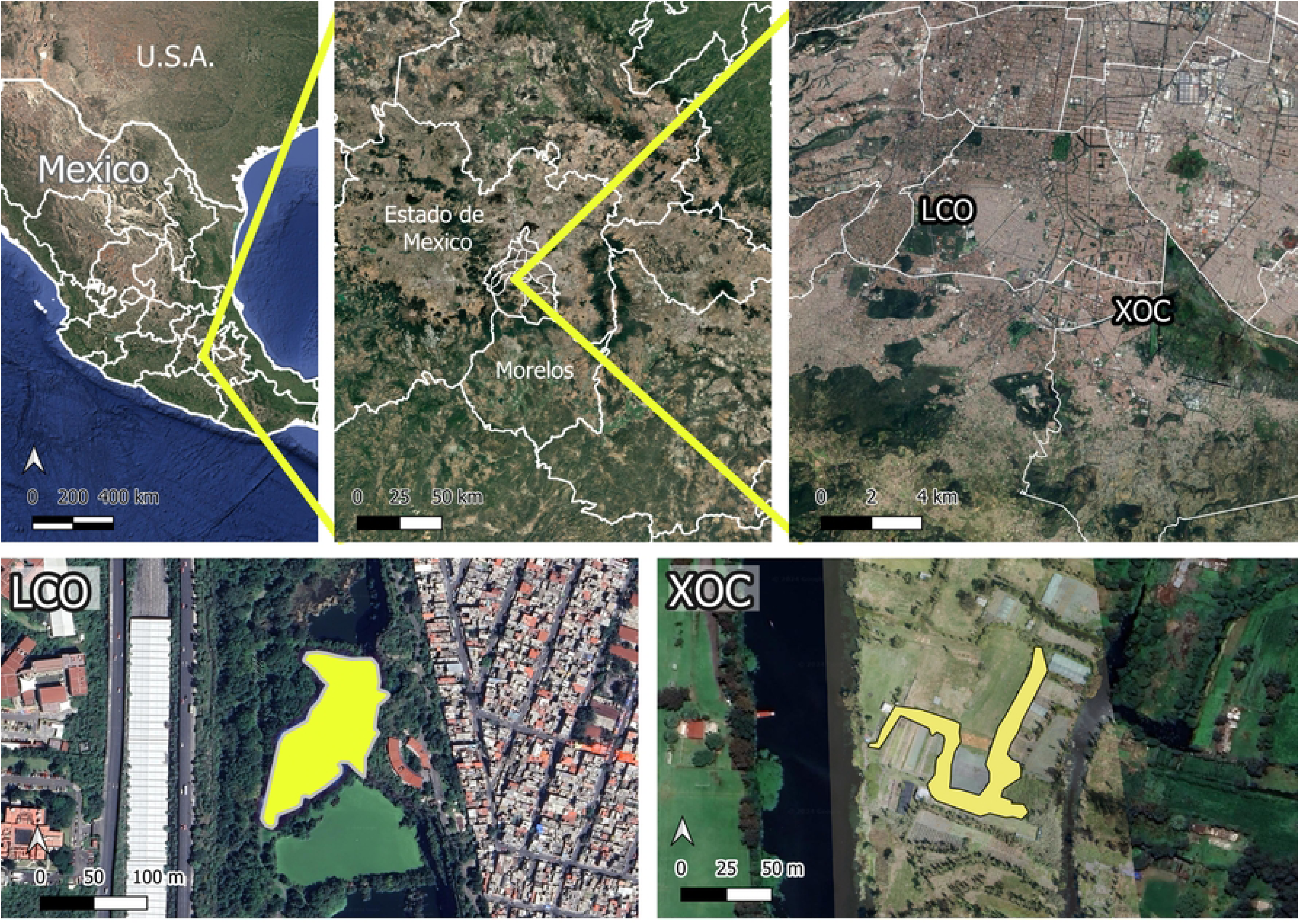
Spatial location of study areas. Overview of the two study areas in southern Mexico City: Xochimilco (XOC) and La Cantera Oriente (LCO). The top panels illustrate the geographic location of the study areas at three distinct spatial scales, while the bottom panels provide close-up views of each wetland. Scale bars represent distances in kilometers (top) and meters (bottom) for reference.

For this study, we selected a 7,059 m² pond within LCO. Previous research has confirmed that this pond provides suitable conditions for axolotl reproduction, including successful egg hatching and larval development into juveniles [39]. Furthermore, in a pilot study, we introduced a radio-tagged, captive-raised male-female pair of axolotls into this pond. Both individuals survived and remained active over a five-week period. Upon recapture, we observed that the male had gained weight and showed no signs of health issues (unpublished data from authors). However, we were unable to recapture the female, as she did not enter our traps.

#### Ethics statement

All axolotls in this study were born and raised in captivity at the Laboratorio de Restauración Ecológica (LRE) of the Instituto de Biología, UNAM, under permit DGVS-PIMVS-CR-IN-1833-CDMX/17. This research complies with Mexican regulations and was conducted under the oversight of the Instituto de Biología’s ethics committee, covering both fieldwork and colony care.

### Study species and transmitters

We studied a total of 18 axolotls, comprising nine females and nine males (S1 Table). The axolotls in the LRE colony are housed without substrate or plants to minimize the risk of disease transmission but are provided with artificial shelters that are routinely sterilized for security and comfort. Their diet includes live food such as artemia, small fish, crayfish, and worms, which are quarantined before feeding and offered every other day, with nutritional supplements administered monthly. Opposite-sex axolotls are kept separately to prevent unintended breeding, and an air-conditioning system maintains the water temperature below 18°C to ensure optimal health.

Axolotls were implanted with SOPI-2038 transmitters from Wildlife Materials, Murphysboro, IL, USA, each programmed with a unique frequency. These transmitters, weighing an average of two grams, constituted about 3% of the axolotls’ body mass, well below the 12% limit deemed acceptable [40]. We used internal transmitters to minimize the risk of skin injuries and to avoid inhibiting natural behaviors such as the use of refugia, predator avoidance, and hunting, which can be affected by external transmitters [34,41,42]

The implantation process began with the axolotls being anesthetized in a 10-minute benzocaine bath (12 ml of benzocaine diluted in 1 L of water). Once sedated, they were weighed and measured. The surgical procedure was performed by Horacio Mena, a veterinarian specialized in axolotls and co-author of this paper. He made a 1-cm longitudinal ventral incision under sterile conditions, carefully inserted the transmitter into the coelomic cavity, and closed the skin with a pattern of spaced stitches, following the administration of analgesics and antibiotics [43]. After surgery, each axolotl was housed individually in a small tank and monitored daily throughout a two-week recovery period. Complete wound healing occurred within a few days without any signs of infection or inflammation.

### Data collection

We utilized a lightweight 3-element YAGI antenna coupled with a Wildlife Materials Inc. (Murphysboro, IL, USA) TRX-48S receiver for radio tracking. Monitoring at both sites was suspended during rainfall or thunderstorms to prevent equipment damage and ensure safety. Initially, we scanned for the unique frequencies of each individual to locate their transmitter signals. Once an axolotl’s signal was detected, team members rowed to the location in LCO or walked in Xochimilco, recording the exact decimal coordinates using a handheld Garmin eTrex 20x GPS unit (Garmin Ltd, Olathe, KA, USA). After completing the final monitoring session, we strategically placed twenty funnel traps in areas where axolotls were most frequently observed. These traps were checked three times daily, every eight hours, over a continuous seven-day period.

On October 26, 2017, we randomly released eight axolotls—four males and four females—at two locations on the north and south sides of the study pond in LCO. From October 27 to December 7, alternating teams of three to four volunteers from a pool of 25, conducted 57 radio-tracking sessions from a small rowboat. We typically scheduled two, and occasionally three, monitoring sessions per day on Mondays, Wednesdays, and Fridays: the first from 9:00 to 12:00 hrs, and the subsequent session(s) from 17:00 to 21:00 hrs. We increased our monitoring efforts in the late afternoons and evenings based on findings from a pilot study, which suggested that axolotls are more active during these times (unpublished data from authors). However, from December 4 to 7— during the final days of this study, radio-tracking sessions were conducted every three hours around the clock to obtain more accurate measurements of distance traveled per hour.

On March 12, 2018, we introduced five male-female pairs of axolotls at five distinct sites within the study Chinampa in Xochimilco, ensuring each site was at least five meters apart. Prior to their release, a small ceremony organized by local Chinamperos took place, during which we offered locally grown flowers to honor the ancient water deities of Xochimilco. From March 12 to April 20, a group of four volunteers from a pool of ten alternated camping at the Chinampa from Monday to Wednesday each week. They conducted radio-tracking of the axolotls on foot at approximately three-hour intervals during this period. However, starting from 18:00 hours on April 16, we increased the frequency of radio-tracking sessions to every hour, continuing until the early morning of April 20.

To document water temperature throughout the day and at the time of each telemetry reading, we placed three HOBO Pendant temperature data loggers (model UA-002-64; Onset Computer Corporation, Bourne, MA, USA) in the LCO study area and eight in the Xochimilco study area. All loggers were positioned at the same depth to ensure consistent measurements, although weather conditions may have influenced water levels. For the temperature analysis, we used the average temperature recorded by all HOBO loggers within each study area at the specific time and hour of each telemetry observation.

### Home range estimation

To estimate the home ranges of individual axolotls in LCO and Xochimilco, we employed the Minimum Convex Polygon (MCP) and enhanced it with the concave k-neighbor method. This advanced approach connects each observation point to its nearest neighbors, forming a boundary that more accurately reflects the actual areas axolotls frequent by excluding extensive, irrelevant territories typically included in traditional MCP estimates [44]. Given that axolotls are strictly aquatic and do not venture outside their water habitats, conventional MCP estimates could overestimate their home ranges.

We conducted a detailed spatial analysis using Kernel Density Estimation (KDE) to identify key habitat areas for individual axolotls and all grouped axolotls independently for both study sites. This method estimates the probability density function of a random variable, allowing us to pinpoint areas of high use from location data. We processed the telemetry data using the sf package [45] to ensure compatibility with spatial analysis tools. We converted it from a tabular format to a spatial features object (sf), enabling direct spatial operations on the dataset.

The KDE model was implemented using the adehabitatHR package in R [46], with the kernelUD function to compute the estimates. We used the reference bandwidth method for smoothing parameter selection. The analysis generated the 50% KDE isopleth, representing the core habitat area where individuals are most frequently found. This isopleth was extracted using the getverticeshr function, providing a spatial representation of the high-density areas for each individual axolotl.

### Data analyses

To compare the biological characteristics and home range sizes between LCO and Xochimilco axolotls, as well as between sexes within each study area, we employed a series of statistical tests. Independent t-tests were used for normally distributed data (mass), while Mann-Whitney U tests were applied to non-normally distributed data (age, length, and home range estimates). Independent t-tests were also used to compare environmental conditions, specifically temperature, between the two study areas.

In addition to these comparisons, we used separate generalized linear mixed models (GLMMs) for each study area to examine the influence of biological, temporal, and environmental factors on axolotl movement, specifically daily and hourly travel distances. Given that the distance data were right-skewed and not normally distributed in both study areas, we applied GLMMs with a Gamma distribution and a log link function for all models. We chose this distribution because it is well-suited for modeling continuous, positively skewed data [47]. Axolotl ID was incorporated as a random effect in all models to account for individual variability.

We assessed collinearity among explanatory variables using Variance Inflation Factors (VIF) to detect multicollinearity in our models. Including both length and mass resulted in VIF values exceeding seven, which indicates multicollinearity. Since mass estimations are more accurate and biologically relevant than length measurements, we excluded length from all analyses. This helped improve model performance and reduce VIF values to below two. We identified the best model for each response variable by comparing AICc values, with model selection performed using the dredge function from the MuMIn package in R [48]. We conducted all statistical analyses using R (v4.4.1) within RStudio 2024.04.2 “Chocolate Cosmos” [49].

#### Distance per hour model

We calculated the hourly travel distance for each axolotl by measuring the distance between consecutive observation points, including those taken up to three hours apart in LCO and up to two hours apart in Xochimilco. For observations that were more or less than one hour apart, we standardized the travel distance to an hourly rate by dividing the observed distance by the time elapsed between points. We then averaged these standardized distances to obtain a consistent measure of hourly travel distance. The explanatory variables in the hourly travel distance models were sex, mass, age, time of day, temperature, and quadratic temperature. We mean-centered both time and temperature.

#### Distance per day model

To calculate the daily total distance traveled by each axolotl, we summed the distances between all consecutive observation points recorded within each 24-hour period. Data from the final days of monitoring in both study areas were excluded because the increased frequency of telemetry readings during this time inflated the estimated daily distances compared to earlier days. The explanatory variables included in the daily distance travel models were sex, mass, age, and number of days since release into the study area.

## Results

We recorded a total of 455 spatial locations in LCO (mean = 56.88 per axolotl; range = 56-57) and 1072 spatial locations in Xochimilco (mean = 107.2 per axolotl; range = 105-110). There were no significant differences in mass, length, or age between axolotls from LCO and Xochimilco, nor between males and females (S2 Table).

At the end of the studies, we successfully recaptured one axolotl in LCO and two in Xochimilco. On January 26, we recaptured a 2.5-year-old female axolotl (A01) in LCO. Initially, she weighed 63.86 grams and measured 20.6 cm. Upon recapture, her weight had increased to 82.2 grams and her length to 21.5 cm, reflecting a gain of 18.34 grams. In Xochimilco, a 2.5-year-old male axolotl (C5), originally weighing 64.3 grams, was recaptured on May 8, showing a 3-gram mass increase and a 3-mm growth in length. Additionally, on May 24, we recaptured a 2.5-year-old female axolotl (A08) with an initial mass of 85.3 grams, although her recapture weight was not recorded. While these recaptures were successful, several other axolotls were detected but evaded capture. Unfortunately, after the study concluded in Xochimilco, we observed a heron capturing an axolotl from the canal. Additionally, local chinamperos reported witnessing another axolotl being taken by a heron, further confirming predation in the area.

### Home Range

The total MCP home range for all eight axolotls in LCO averaged 2,747 m² (range: 548–4,404 m²) (Fig 2), while ten axolotls in Xochimilco showed a smaller mean of 382 m² (range: 175-697 m²) (Fig 3). This difference was statistically significant (p = 0.00018), indicating that axolotls in LCO utilize a much larger area than those in Xochimilco. Similarly, the KDE 50% areas for the eight axolotls in LCO averaged 1,640 m² (range: 211–3,673 m²), compared to a smaller mean of 204 m² (range: 35-455 m²) for the ten axolotls in Xochimilco. This difference was also statistically significant (p = 0.0014). We found no significant differences in MCP or KDE 50% core areas between sexes in either study area (LCO and Xochimilco), additionally, there were no significant correlations between MCP or KDE 50% sizes and age in either location (Table S2).

**Figure 2.**
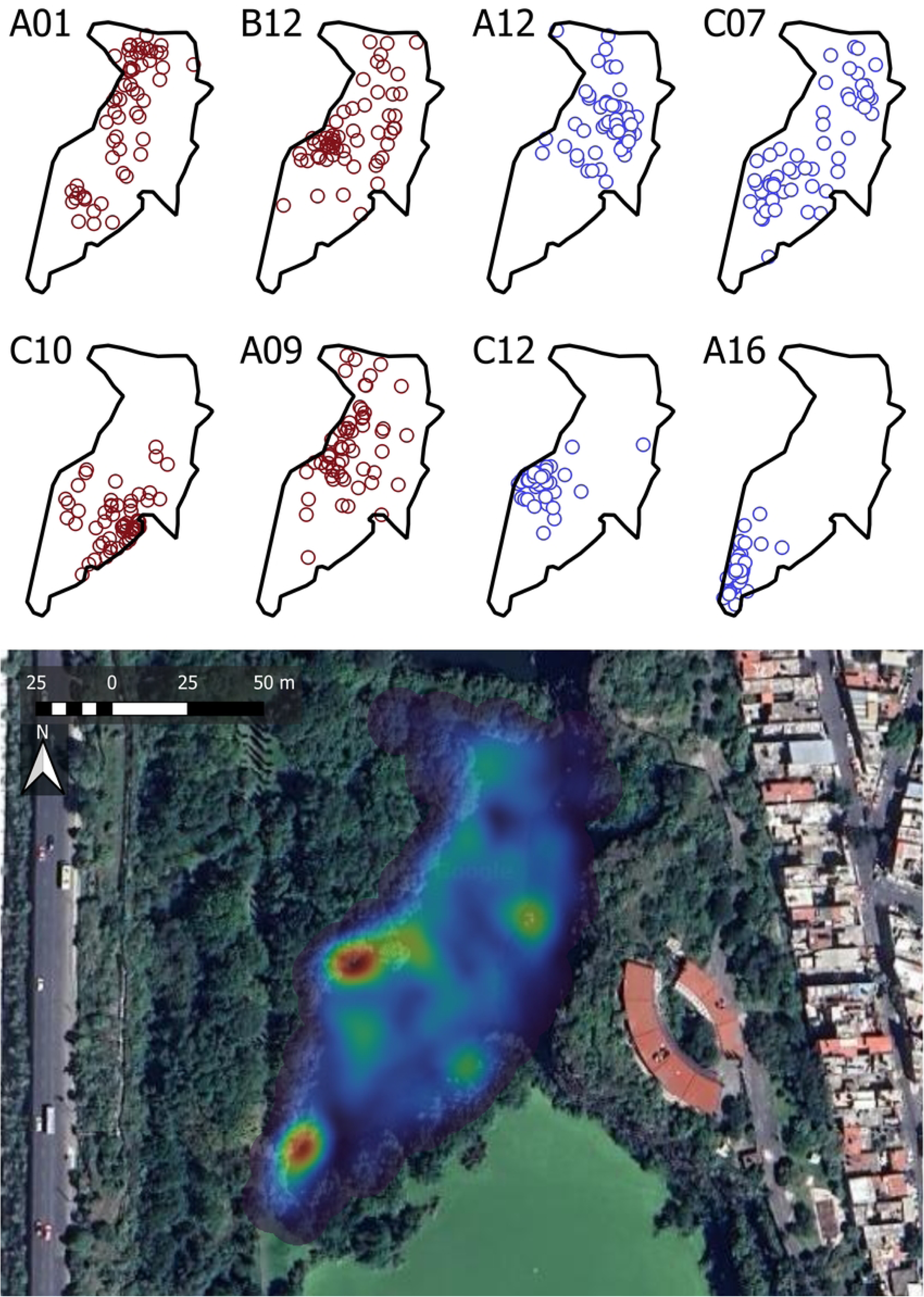
Spatial distribution and habitat use in LCO. Top panel: distribution of individual axolotls, separated by sex, with males shown in blue (IDs: A12, C07, C12, A16) and females in red (IDs: A01, B12, C10, A09). Bottom panel: Kernel Density Estimation (KDE) heatmap illustrating habitat use intensity, with warmer colors indicating areas of higher use.

**Figure 3.**
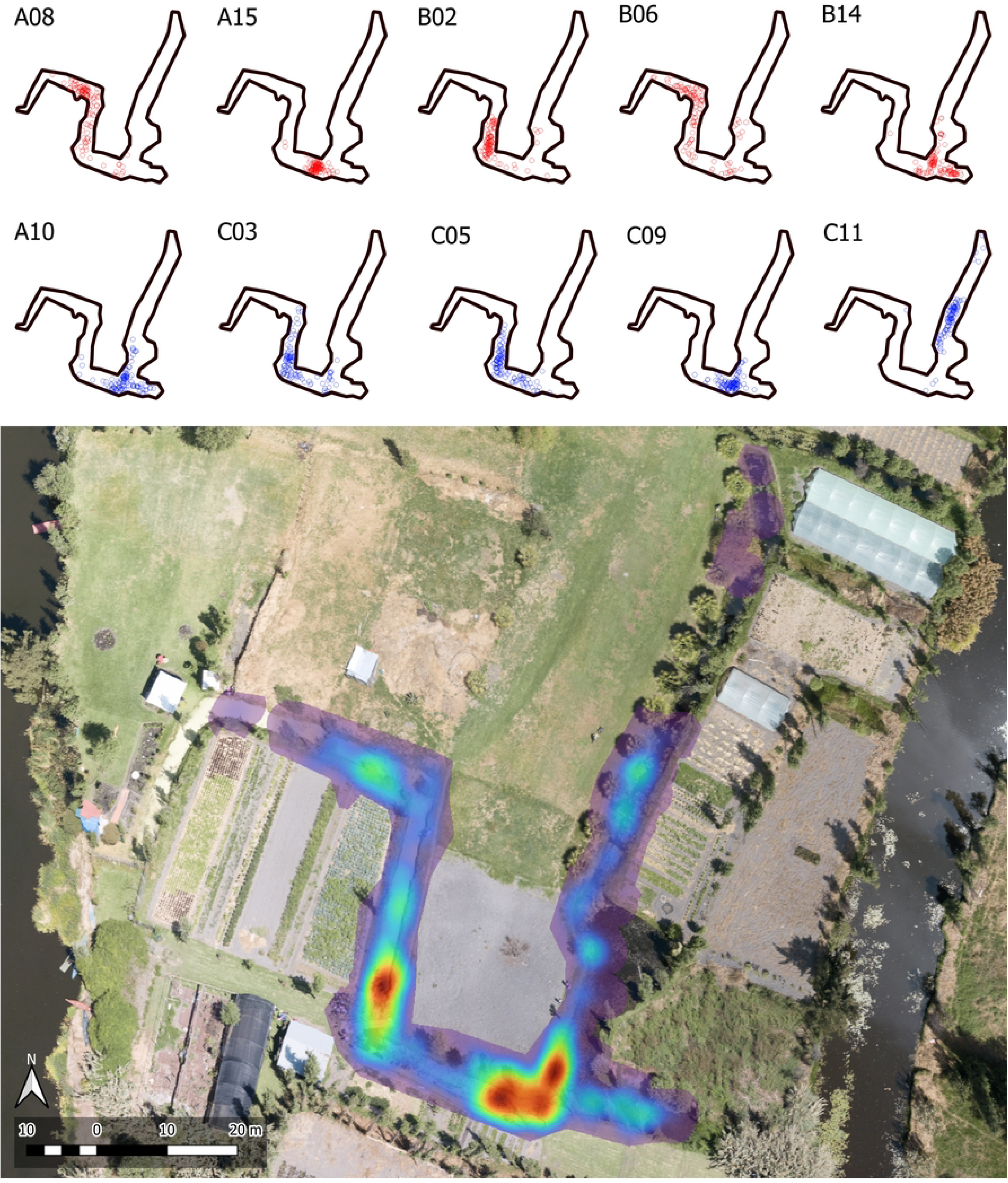
Spatial distribution and habitat use in Xochimilco. Top panel: distribution of individual axolotls separated by sex, with males shown in blue (IDs: A10, C03, C05, C09, C11) and females in red (IDs: A08, A15, B02, B06, B14). Bottom panel: Kernel Density Estimation (KDE) heatmap representing habitat use intensity, with warmer colors indicating areas of higher use.

### Distance per Hour

The mean water temperature at LCO, based on readings corresponding to telemetry data collection times, was 15.92°C (range: 14.58-17.73; n = 222), compared to 16.66°C at Xochimilco (range: 14.9-18.73; n = 439). An independent t-test confirmed that LCO was significantly cooler than Xochimilco (p < 0.001). When considering all temperature data recorded by the HOBO devices, more extreme values were observed. The overall mean water temperature at LCO, using all recorded data, was 16.34°C (range: 12.11-30.76; n = 8,307), while at Xochimilco, it was 16.52°C (range: 12.01-37.71; n = 14,744). The best-fitting model for LCO revealed a significant quadratic relationship between distance traveled per hour and water temperature (p < 0.001), with movement increasing with temperature up to a peak, then decreasing at higher temperatures (Fig. 4). In Xochimilco, we found a similar significant quadratic relationship (p < 0.001) that mirrored the pattern observed in LCO (Fig. 4). In addition, the LCO model revealed a significant effect of sex on distance traveled per hour, with males covering less distance compared to females (p = 0.032). While the time of day also had a significant impact on movement (p = 0.019), indicating an increase in activity levels during the later hours of the day. Neither sex nor time of day were significant in the Xochimilco.

**Figure 4.**
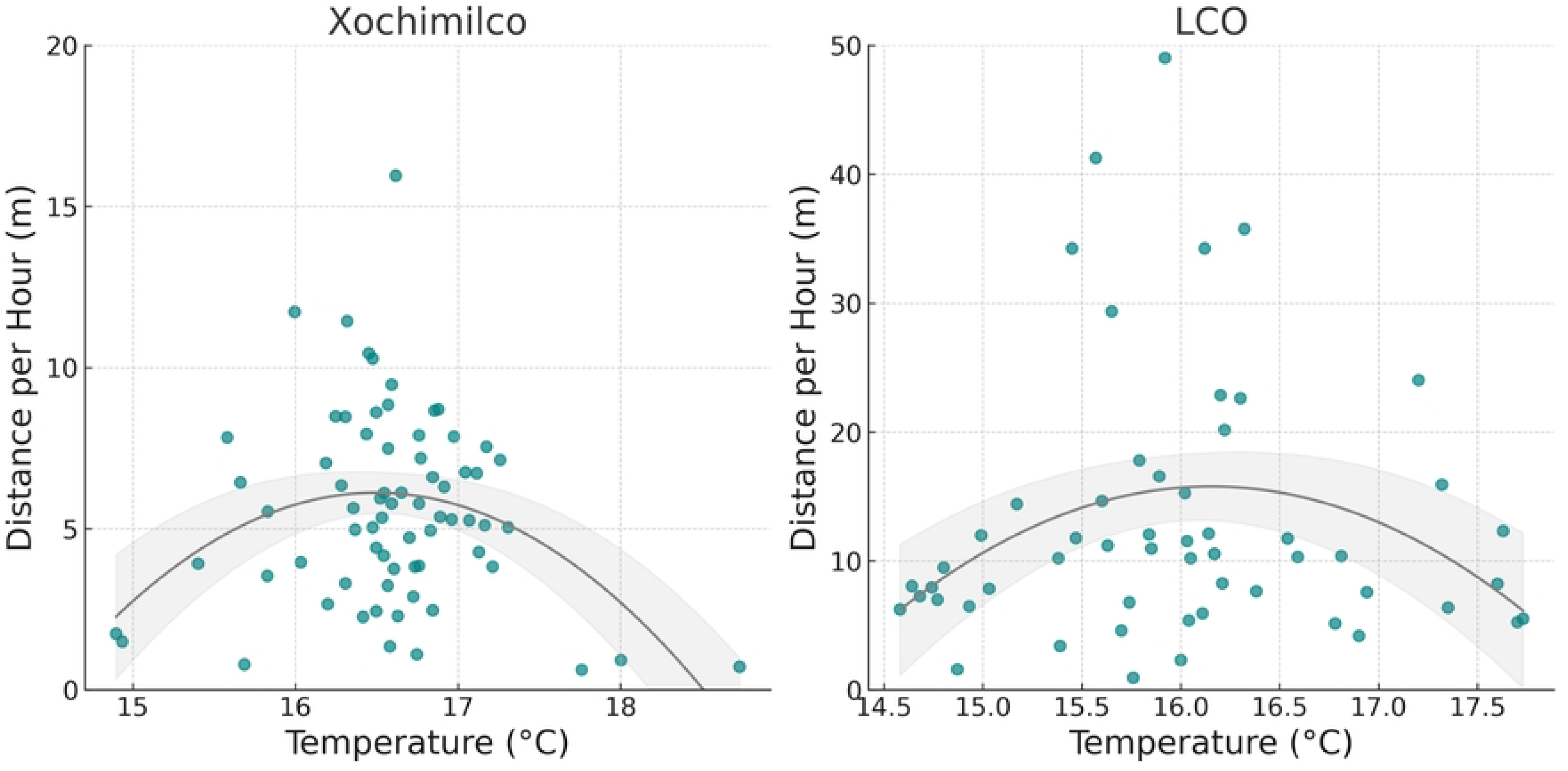
Relationship between distance traveled per hour and water temperature. The left panel shows data from Xochimilco, and the right panel shows data from LCO. Data points represent the average distance traveled per hour for each unique temperature value, plotted in teal. The grey line represents a quadratic trend line with corresponding confidence intervals.

### Distance per Day

We analyzed the daily distances traveled by axolotls at two different sites. In LCO, the mean distance traveled per day was 86.3 meters (range: 6.79-301.54 meters). In contrast, axolotls in Xochimilco traveled a mean distance of 51.9 meters per day (range: 2.39-174.72 meters). Axolotls in LCO traveled statistically larger distances than those in Xochimilco (U = 6772.5, p < 0.001).

The distance analysis for LCO showed that the number of days since release had a significant negative effect on the daily distance traveled by axolotls (p = 0.0177; Fig 5). This suggests that the daily distance traveled by axolotls decreased over time. Mass was marginally significant (p = 0.055), indicating a trend where larger axolotls tend to travel shorter distances each day. Neither sex, age, nor their interactions were included in the best model, as they did not significantly improve model fit. For the Xochimilco axolotls, the number of days since release did not have a significant effect on daily distance traveled. However, age had a significant negative effect (p = 0.0077; Fig 6), indicating that older axolotls traveled shorter distances. Sex, mass, and interactions between covariates were not included in the final model as they did not significantly improve model fit.

**Figure 5.**
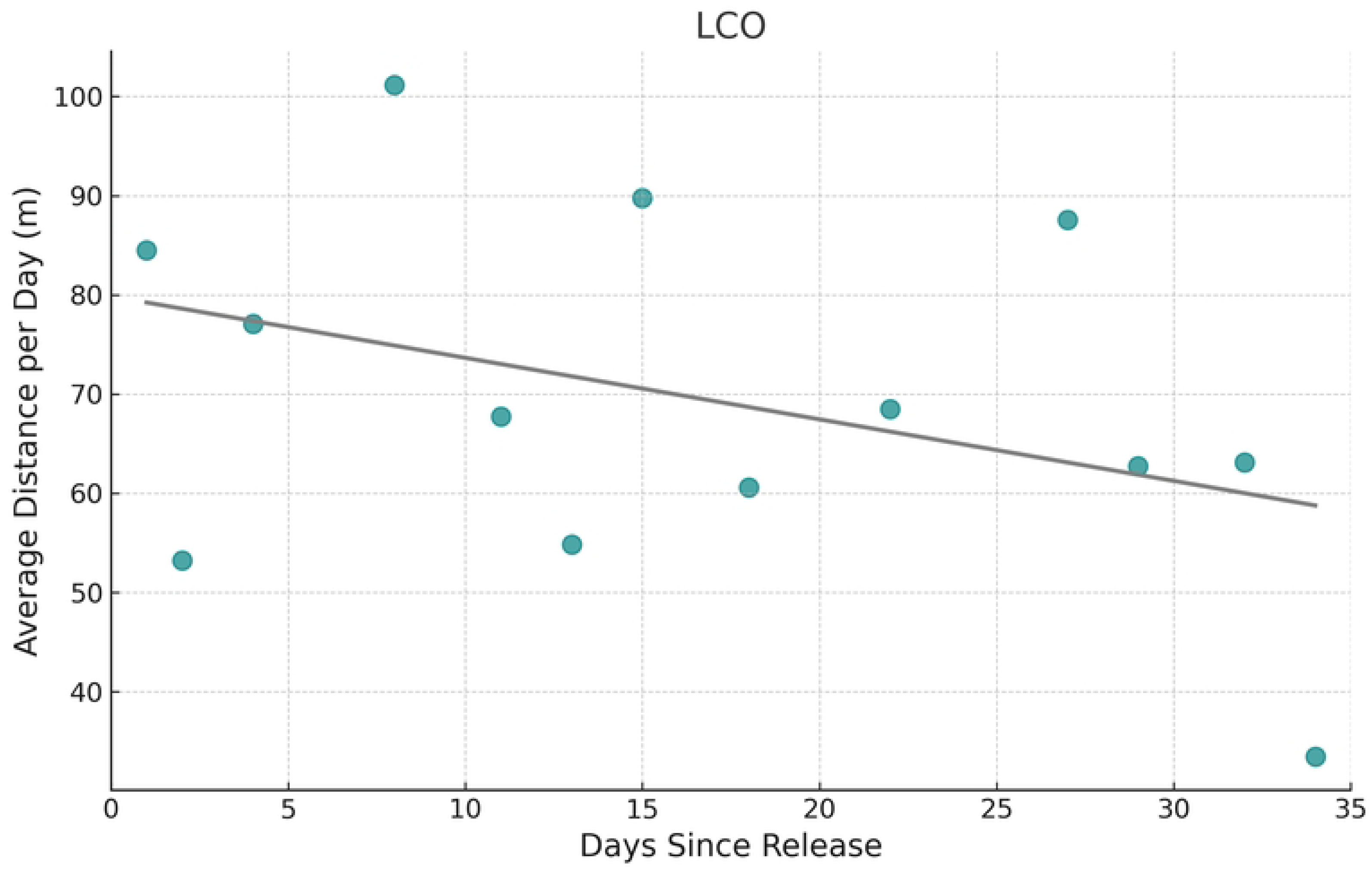
Effect of time since release on daily travel distance in LCO axolotls. Relationship between the number of days since release into the study area and the average distance traveled per day by axolotls.

**Figure 6.**
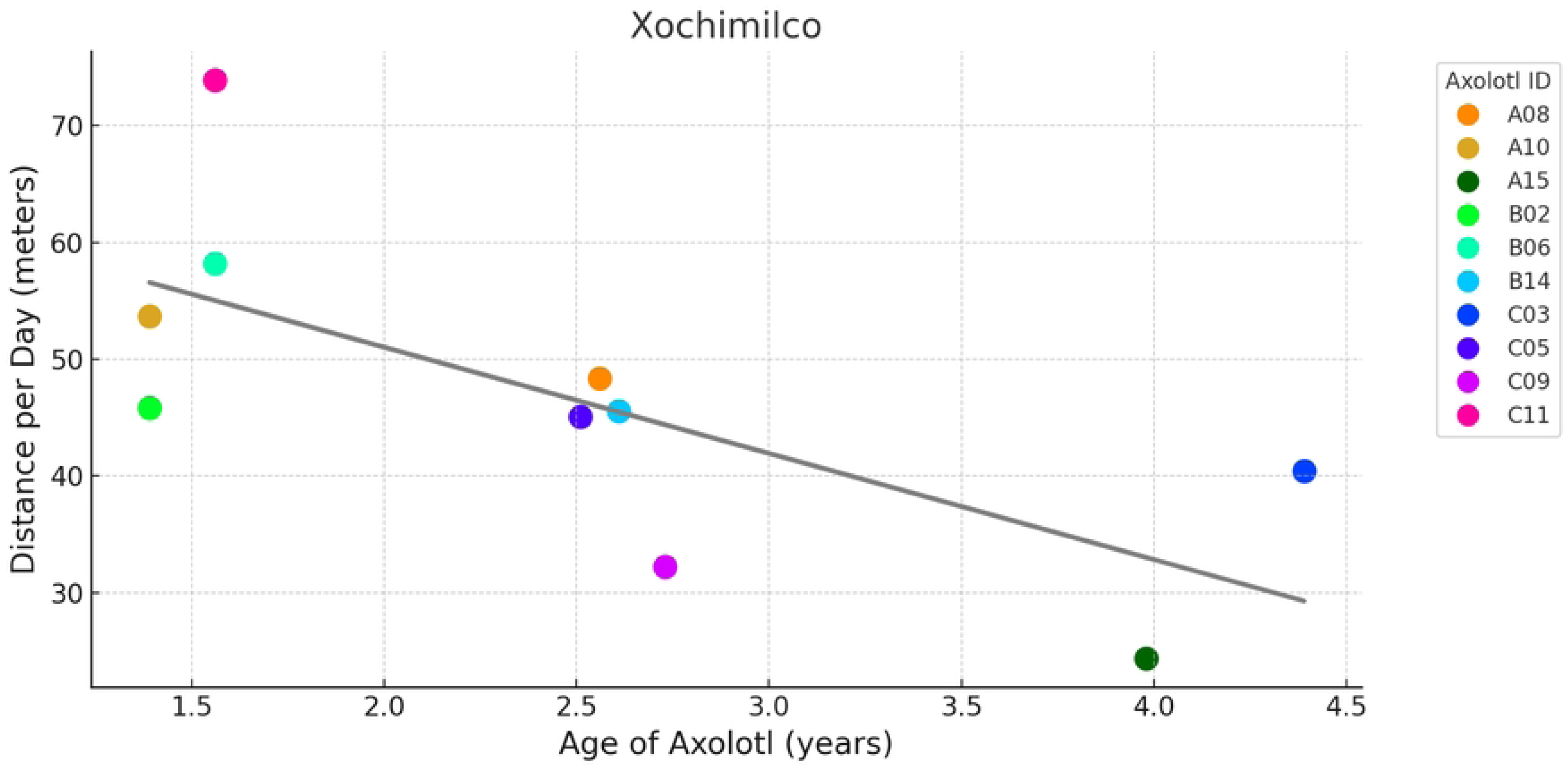
Influence of age on daily travel distance in Xochimilco axolotls. Each circle represents the average distance traveled per day by an individual axolotl, color-coded uniquely for each axolotl ID. Jitter was applied to distinguish overlapping points for axolotls B14 and C05.The grey line represents the linear trend of the data.

## Discussion

We used VHF telemetry to explore the movement behaviors and home range sizes of captive-bred axolotls released in both an artificial wetland (LCO) and their native habitat (Xochimilco). Axolotls in both study areas survived the approximately 40-day monitoring period. Of the three recaptured individuals, two that were reweighed—one from each study area—showed weight gain, suggesting successful foraging in both locations. However, shortly after the study in Xochimilco concluded, we observed a heron capturing an axolotl from the canal, and chinamperos later reported a second axolotl falling prey to a heron, indicating elevated predation in this location.

Axolotls in LCO and Xochimilco were not randomly distributed but instead showed a preference for certain microhabitats, as evidenced by high densities of observations in certain areas and minimal to no observations in others (Figs 2 and 3). In Xochimilco, axolotls notably avoided the westernmost section of the canal, where dense mats of floating duckweed (*Lemna gibba*) covered the water surface. Similarly, a study on spotted salamanders (*Ambystoma maculatum*) found fewer adults, egg masses, and larvae in pools with significantly more duckweed cover compared to those with less [50]. However, neither our study nor the other confirmed a causal relationship between duckweed presence and salamander distribution. Habitat preferences have been fairly documented in aquatic salamanders. Hellbenders (*Cryptobranchus alleganiensis*), for example, often hide under large, flat rocks and select microhabitats with gravel substrate over others [51,52]. Giant salamanders (*Andrias davidianus*) seek crevices that offer protection against predators while providing easy access to the surface [53]. Reintroduced axolotls (*A. mexicanum*), likewise, demonstrate a preference for aquatic vegetation, such as *Myriophyllum aquaticum* and *Eichhornia crassipes*, during daytime, likely as a strategy for predator avoidance during hours in which they are less active [37].

Xochimilco axolotls exhibited smaller home range sizes compared to those in LCO. This difference can likely be attributed to the considerably larger water area of the LCO pond, which is over 10 times the size of the Xochimilco canal. While we did not find direct evidence supporting this relationship in salamanders, water body size has been shown to be a critical factor in determining the home range sizes of freshwater fish [54,55]. Nevertheless, other factors might also influence movement patterns. For instance, research on hellbenders, another species of aquatic salamanders, suggests that habitat features, such as substrate and shelter availability, can significantly influence movement patterns and home range sizes [31,56]. In these studies, hellbenders released into areas with larger, more connected boulder formations tended to travel shorter distances, maintain more compact home ranges, and exhibit greater site fidelity compared to those placed in environments with more scattered boulders. Furthermore, another study found that translocated hellbenders moved greater distances and established larger home ranges compared to resident individuals, likely due to exploratory behavior, habitat suitability, and predation risks [57].

In LCO, axolotls showed a significant decrease in daily distance traveled as the number of days since release increased, suggesting that individuals reduced their movement over time as they became more settled in their environment. In contrast, axolotls in Xochimilco did not exhibit this pattern, likely because the smaller and simpler habitat provided quicker access to necessary resources. This finding from LCO is consistent with studies on hellbenders, which typically exhibit a short exploration phase before achieving stable settlement and high site fidelity within three weeks post-release. Similarly, captive-reared Chinese giant salamanders reintroduced to the wild were observed to settle in a single location within 10 days post-release [33]. Exploratory behavior is essential after translocations and reintroductions, as it allows animals to locate resources like food and shelter, while also assessing predator threats [58,59]. However, this behavior can also be costly, as it requires energy and heightens the risk of predation [60,61]. The larger, more complex LCO pond likely required axolotls to engage in a longer exploratory phase, as they navigated the environment to find optimal foraging and shelter sites, further contributing to their larger home ranges.

The distance traveled per hour by LCO axolotls showed a significant positive relationship with the time of day, with activity levels increasing in the afternoon and peaking around 9:00 pm. This rise in nighttime activity could be influenced by a combination of factors, including nocturnal foraging behavior, as observed in hellbenders [62], and predator avoidance, as documented in Northwestern salamanders (*Ambystoma gracile*) [63]. In a pilot study with two axolotls translocated into the LCO pond and monitored using VIF equipment, we observed that after an initial exploratory period of roughly one month, both individuals exhibited a clear shift in spatial use: they spent the daytime hours, when they were less active, in the northern part of the pond, and the nighttime hours in the southern section, where we suspect food availability was higher (unpublished data by the authors; S3 Fig). Although the current study did not reveal a distinct day-night spatial pattern, there were concentrations of daytime observations in northern areas, while nighttime movements were more frequent in the central and southern regions. In contrast, our current study in Xochimilco did not show a significant relationship between time of day and activity. However, in a previous experimental study conducted in Xochimilco, where we created distinct vegetated and non-vegetated quadrants, axolotls were more active at night and spent more time in non-vegetated areas [37]. The lack of a significant relationship in the current study could be due to the absence of environmental manipulation, leading to a more natural and homogenous distribution of vegetation compared to the experimental canal.

The relationship between axolotl movement, measured as distance traveled per hour, and temperature followed a nonlinear trend in both study areas. Axolotl movement increased with temperature up to a certain point, peaking at around 16– 17°C in Xochimilco and 15.5–16.5°C in LCO and, beyond these optimal ranges, movement declined as temperatures continued to rise (Fig 6). This suggests that axolotls are most active within a narrow temperature window, and higher or lower temperatures may reduce movement, potentially reflecting physiological constraints. Previous research on amphibians has shown that temperature significantly influences their movement patterns, as they rely on external heat to regulate their metabolism and activity, particularly in response to environmental conditions [64]. Several studies have documented observed and preferred temperatures of salamanders in both field and laboratory conditions [65]. For example, *A. mexicanum* in laboratory settings selected temperatures between 16°C to 18°C [66], which resembles the optimal movement temperatures observed in our study. Similarly, mountain stream salamanders (*Ambystoma altamirani*) from Central Mexico were found at temperatures ranging from 16.42°C to 16.98°C [67]. However, several other *Ambystoma* species from Noth America have been associated with considerably higher temperatures, ranging from 24.8 °C to 34.6 °C [65].

On average, water temperatures in Xochimilco during March and April were significantly warmer than those in LCO from late October to early December. Interestingly, the minimum (12°C) and maximum (38°C) temperatures recorded in Xochimilco matched those reported in a single, shallow sample site from a 2003-2004 study [68]. However, our observed mean water temperature was lower, at 16.5°C, compared to the 20.5°C estimated from data across all sites in their study. It is important to note that our study did not include the warmest months of the year, whereas García et al.’s study collected data year-round. Given that we recorded identical extreme temperatures without sampling the warmest months, it is possible that maximum temperatures during the summer are now even higher, especially in shallow areas, potentially due to climate change’s impact on rising temperatures. From a conservation perspective, this highlights the potential importance of LCO as a complementary long-term strategy for axolotl conservation, especially if it remains cooler than Xochimilco in the future. Additionally, the conditions at LCO may better reflect Xochimilco’s historical state, before the Aztecs transformed much of it into chinampas, a system of artificial islands used for agriculture that altered the natural water flow and ecosystem. Since axolotls evolved in the natural lake of pre-Aztec Xochimilco, they are likely better adapted to these historical conditions, which LCO may now better resemble.

At the individual level, our results show that age and sex significantly influenced axolotl movement measured by hourly and daily traveled distance. In Xochimilco, older axolotls traveled shorter daily distances compared to younger ones (Fig 5), while in LCO, males covered less hourly distance than females. However, we found no relationship between individual traits and home range sizes. While much research has focused on the influence of environmental factors on movement patterns like dispersal in amphibians, there remains limited empirical evidence regarding the role of individual traits, such as age, sex, and body size [32]. Juvenile Chinese giant salamanders have been observed to disperse farther than adults, likely moving away from their natal population [33], which aligns with our finding that younger axolotls were more mobile. Similarly, dispersal distance in red-backed salamanders (*Plethodon cinereus*) was significantly greater in juveniles than in adults [69], further supporting the idea that juveniles tend to be more mobile during early life stages across different species. This variation in movement patterns seen between juvenile and adult salamanders could also be attributed to territorial behavior in adults, as adults are known to establish and defend territories [33]. Though environmental factors may influence space use differently based on age and sex, identifying the sex of salamanders can be particularly challenging [70]. Additionally, studies that include age often categorize it into distinct life stages (larvae, juvenile, and adult), likely because they lack precise age estimates, rather than treating it as a continuous variable as we did in this study.

Our study demonstrates that captive-bred axolotls can survive and successfully forage in both their native habitat of Xochimilco and an artificial wetland environment like LCO. The observation of weight gain in recaptured individuals from both locations indicates effective foraging, although elevated predation risks in Xochimilco, evidenced by heron attacks, remain a significant concern. One of the main causes of failure in reintroduction and translocation programs is the high mortality due to predation that occurs after release [71,72]. Animals born and raised in captivity do not learn to recognize and respond appropriately to predators and may completely lose their anti-predator behaviors, or these behaviors may become less efficient compared to those of wild animals [73]. Amphibians may be particularly at risk, as their small size makes them highly susceptible to predation by a wide range of predators, including reptiles, birds, and mammals [64].

The use of animal behavior as a tool for conservation has shown promise in improving survival rates of threatened species reintroduced into the wild [74]. ‘Pre-release training’ can play a crucial role in mitigating predation risks and boosting the success of reintroductions. Research on mammals, birds, and fish indicates that captive-born individuals can learn to recognize and respond to predators if given the appropriate training before release [73]. For example, Puerto Rican parrots (*Amazona vittata*) exposed to cues simulating predation risk from hawks were less likely to be preyed upon by raptors compared to those that did not receive training [75]. Similarly, amphibian embryos and larvae can be trained with chemical stimuli to recognize predators [76]. Notably, responses to these stimuli may vary with age; for example, larvae of the giant American salamander (*Cryptobranchus alleganiensis*) trained to respond to trout cues remained still at around 4 months old but fled at around 6 months old [77].

Applying pre-release training to axolotls could reduce predation and enhance survival post-release. Temporary shelters could serve as pre-release enclosures where axolotls are exposed to visual and olfactory predator cues, mimicking real-life threats to help condition their anti-predator behaviors before they are released into the wild. In a previous study, shelters were successfully used in LCO where adult axolotls reproduced, and their eggs hatched into larvae, which then grew into juveniles [39]. This approach could enhance the success of reintroduction programs by offering axolotls a safer, semi-natural environment during the critical early stages of development, thereby strengthening their behavioral responses to threats and increasing their chances of long-term survival in the wild.

Our findings provide valuable insights into the movement ecology and habitat use of axolotls, informing conservation efforts aimed at their reintroduction and habitat management. By demonstrating the ability of captive-bred axolotls to survive and forage in both native and artificial environments, we reinforce the potential of artificial wetlands like LCO to serve as supplementary habitats for this critically endangered species. The elevated predation risks observed in Xochimilco highlight the necessity of implementing strategies such as pre-release predator training to enhance survival rates. Additionally, while our study did not directly compare temperatures simultaneously across sites, the observed differences suggest that LCO may offer a cooler, more stable environment. This could be particularly advantageous given the expected impacts of climate change on axolotl habitats. Continued monitoring and comparison of environmental variables such as temperature and water quality will be crucial in refining the use of LCO as a complementary conservation tool alongside Xochimilco. Future research should focus on refining pre-release training methods, assessing long-term survival and reproduction post-release, and comparing key environmental factors between LCO and Xochimilco to refine conservation strategies. Overall, our study contributes to a better understanding of axolotl ecology and offers practical approaches to improve conservation outcomes for this iconic species.

## Acknowledgements

We are grateful to the many field assistants and volunteers who contributed their time and effort to this study in Xochimilco and La Cantera Oriente. We especially acknowledge Carlos Uriel Sumano Arias, Esmeralda Benitez Sandoval, Omar Jiménez, Cyntia Giselle Pensado Ortega, Araceli Mejía Barrio, Pedro Alberto Cabrera Castillo, Lourdes Itzel Murillo Reyes, Ana Yunuen Aguilera León, Allison Lory Bettencourt, Diana Laura Vázquez Mendoza, and Fabiola Jocelyn Real Reyes, who assisted in the field across one or both study areas. The contributions of additional volunteers, too numerous to list here individually, are also deeply appreciated.

## Supporting information

**S1 Table. Individual traits of LCO and Xochimilco axolotls.** Summary of individual characteristics of axolotls studied across two different study areas, LCO and Xochimilco. The table lists the identification code (ID), sex, age (in years), body mass (in grams), and length (in centimeters) of each axolotl.

**S2 Table. Biological and spatial parameter comparison of axolotls by location and sex.** Comparison of individual traits (mass, length, and age) and home range (MCP and KDE 50%) between axolotls from LCO and Xochimilco, and between sexes within each location. Values are presented as means with ranges in parentheses. Significant p-values are highlighted in bold.

**S3 Fig. Spatial distribution of axolotl observation points in LCO**. The pilot study is displayed on the left, while the current study is on the right. Yellow points represent axolotl locations observed during the day, and purple points represent locations observed at night.

